# SARS-CoV-2 Infects Syncytiotrophoblast and Activates Inflammatory Responses in the Placenta

**DOI:** 10.1101/2021.06.01.446676

**Authors:** Lissenya B. Argueta, Lauretta A. Lacko, Yaron Bram, Takuya Tada, Lucia Carrau, Tuo Zhang, Skyler Uhl, Brienne C. Lubor, Vasuretha Chandar, Cristianel Gil, Wei Zhang, Brittany Dodson, Jeroen Bastiaans, Malavika Prabhu, Christine M. Salvatore, Yawei J. Yang, Rebecca N. Baergen, Benjamin R. tenOever, Nathaniel R. Landau, Shuibing Chen, Robert E. Schwartz, Heidi Stuhlmann

## Abstract

SARS-CoV-2 infection during pregnancy leads to an increased risk of adverse pregnancy outcomes. Although the placenta itself can be a target of virus infection, most neonates are virus free and are born healthy or recover quickly. Here, we investigated the impact of SARS-CoV-2 infection on the placenta from a cohort of women who were infected late during pregnancy and had tested nasal swab positive for SARS-CoV-2 by qRT-PCR at delivery. SARS-CoV-2 genomic and subgenomic RNA was detected in 23 out of 54 placentas. Two placentas with high virus content were obtained from mothers who presented with severe COVID-19 and whose pregnancies resulted in adverse outcomes for the fetuses, including intrauterine fetal demise and a preterm delivered baby still in newborn intensive care. Examination of the placental samples with high virus content showed efficient SARS-CoV-2 infection, using RNA in situ hybridization to detect genomic and replicating viral RNA, and immunohistochemistry to detect SARS-CoV-2 nucleocapsid protein. Infection was restricted to syncytiotrophoblast cells that envelope the fetal chorionic villi and are in direct contact with maternal blood. The infected placentas displayed massive infiltration of maternal immune cells including macrophages into intervillous spaces, potentially contributing to inflammation of the tissue. *Ex vivo* infection of placental cultures with SARS-CoV-2 or with SARS-CoV-2 spike (S) protein pseudotyped lentivirus targeted mostly syncytiotrophoblast and in rare events endothelial cells. Infection was reduced by using blocking antibodies against ACE2 and against Neuropilin 1, suggesting that SARS-CoV-2 may utilize alternative receptors for entry into placental cells.

## Introduction

The global pandemic resulting from the novel coronavirus, Severe acute respiratory syndrome coronavirus 2 (SARS-CoV-2), has already taken a devastating toll, with over 175 million total cases and more than 3.8 million deaths worldwide. SARS-CoV-2 which causes Coronavirus Disease 2019 (COVID-19) has significant clinical variability. In severe cases SARS-CoV-2 causes a respiratory illness, whose defining features are an imbalanced inflammatory host response, reduced innate antiviral defenses and an inflammatory “cytokine storm”, endothelial damage, coagulopathies and thrombosis in several tissues from infected patients (Blanco-Melo et al. 2020).

To date, our understanding of how SARS-CoV-2 infection impacts pregnancy, including the health of COVID-19 positive mothers and their babies, remains incomplete. Pregnant women with symptomatic COVID-19 infections are more likely to be admitted to the intensive care unit (ICU), and have statistically higher maternal death rates when compared to non-pregnant infected women (Zambrano et al. 2020). While preterm deliveries occur more often in women with suspected or confirmed SARS-CoV-2 infection, no increase in stillbirth or early neonatal death was found (Mullins et al. 2021). Prospective and retrospective studies showed that pregnant women infected with SARS-CoV-2 are at increased risk of adverse events, including higher rates of cesarean section and increased post-partum complications (Woodworth et al. 2020; Prabhu et al. 2020; Marín Gabriel et al. 2020). While vertical transmission of SARS-CoV-2 from mother to fetus has been reported in a few cases (Hecht et al. 2020a; Vivanti et al. 2020; Taglauer et al. 2020; Facchetti et al. 2020; Woodworth et al. 2020; Hecht et al. 2020b; Alamar et al. 2020), most studies did not detect viral transmission (Penfield et al. 2020; Baergen and Heller 2020; Prabhu et al. 2020; Salvatore et al. 2020; Edlow et al. 2020; Schwartz 2020; Della Gatta et al. 2020; Kimberlin and Stagno 2020).

Several studies have detected SARS-CoV-2 infection of the placentas from women who tested positive for the virus at, or prior to, delivery. In some cases, the placenta displayed signs of inflammation. These placentas were found to have increased vascular malperfusion indicative of thrombi in fetal vessels (Baergen and Heller 2020; Vivanti et al. 2020; Prabhu et al. 2020; Shanes et al. 2020) and infiltration of maternal immune cells (Hosier et al. 2020; Facchetti et al. 2020; Debelenko et al. 2021; Garrido-Pontnou et al. 2021; Lu-Culligan et al. 2021; Morotti et al. 2021; Schwartz et al. 2021). Whether inflammation results from virus infection of the mother or direct infection of the placenta remains unresolved, as this may depend on the gestational age of the fetus. Virus infection is known to impair placental function. Virus-associated inflammation during pregnancy can result in chronic cardiovascular disease, diabetes and obesity later in life (Burton, Fowden, and Thornburg 2016). Little is known however about the effect of SARS-CoV-2 on placental function.

SARS-CoV-2 utilizes ACE2 (Angiotensin-converting enzyme 2) as the primary receptor (Hoffmann et al. 2020), and Neuropilin-1 (NRP1) as a coreceptor (Cantuti-Castelvetri et al. 2020; Daly et al. 2020), in concert with the two proteinases TMPRSS2 (Transmembrane protease serine 2) (Hoffmann et al. 2020) and CTSL (Ou et al. 2020) and the pro-protein convertase furin (Shang et al. 2020), amongst others (Wei et al. 2021; Daniloski et al. 2021; Wang et al. 2021; Schneider et al. 2021) for cell entry. All of the viral entry receptors are expressed at significant levels in first and second trimester placentas. However, at term they are expressed at lower levels at the maternal-fetal interface, including the placenta and the chorioamniotic membranes (Pique-Regi et al. 2020; Li et al. 2020; Singh, Bansal, and Feschotte 2020; Taglauer et al. 2020; Lu-Culligan et al. 2021; Baston-Buest et al. 2011). Whether alternative entry mechanisms are exploited by SARS-CoV-2 in the placenta is not known.

In this study we were interested to understand the impact of SARS-CoV-2 infection late in pregnancy on placental function. Using a cohort of 54 women who tested positive for SARS-CoV-2 at the time of delivery, we report on placental infections, placental pathologies and *in vivo* inflammatory responses to infection. Furthermore, we present *in situ* studies of infected placentas as well as *ex vivo* placental explant cultures that investigate susceptibility of placental cells to SARS-CoV-2 infection.

## Results

### Clinical presentations of SARS-CoV-2 positive mothers, fetal outcomes and placental pathologies

All placental samples in the present study were provided by the Department of Pathology and Laboratory Medicine at Weill Cornell Medicine. A cohort of 54 women who were identified as positive for SARS-CoV-2 by RT-PCR from nasopharyngeal swabs at the time of admission for delivery at NY Presbyterian Hospital-Weill Cornell was included (P1–P54). As controls, a cohort of 5 women who tested negative for SARS-CoV-2 (C1-C5), and a cohort of 5 SARS-CoV-2 negative women who presented various placental inflammatory pathologies (I1-I5) were also included in the study (Table 1).

**Table 1.** Clinical presentations of SARS-CoV-2 positive mothers, fetal outcomes and placental pathologies. Overview of 65 Patients included in study. 55 COVID-positive, 10 COVID-negative. Abbreviations: FVM: Fetal Vascular Malperfusion, MVM: Maternal Vascular Malperfusion, DFM: Decreased Fetal Movement, MCI: Massive Chronic Intervillositis, MFI: Maternal Floor Infarction, CHI: Chronic Histiocytic Intervillositis, IUFD: Intra-Uterine Fetal Demise, T2D: Type 2 Diabetes, Mec: Meconium, IVT: Intervillous Thrombi, VUE: Villositis of Unknown Etiology/Chronic Villositis, ACA: Acute Chorioamniotis, IAI: Intra-Amniotic Infection/Chorioamnioitis, HTN: Hypertension, IUGR: Intra-Uterine Growth Restriction, GDM: Gestational Diabetes Mellitus, PPROM: Preterm Premature Rupture of Membranes, PTL: Preterm Labor, PAPP-A: Pregnancy-associated Plasma Protein A, UCTD: Undifferentiated Connective Tissue Disorder, HCV: Hepatitis C Virus, ITP: Immune Thrombocytopenic Purpura. IUGR: intrauterine growth restriction. ICP: intrahepatic cholestasis of pregnancy. Gray Shaded Rows = Fetal Demise/NICU admission. DFM: Decreased Fetal Movement, MCI: Massive Chronic Intervillositis, MFI: Maternal Floor Infarction, CHI: Chronic Histiocytic Intervillositis, IUFD: Intra-Uterine Fetal Demise, T2D: Type 2 Diabetes, Mec: Meconium, IVT: Intervillous Thrombi, VUE: Villositis of Unknown Etiology/ Chronic Villositis, ICP: Intrahepatic Cholestasis of Pregnancy, GBS: Group B Streptococcus+, TOLAC: Trial of Labor After Cesarean, ACA: Acute Chorioamnionitis, BMI: Body Mass Index, TAB: Therapeutic Abortion, IDA: Iron Deficiency Anemia, IAI: Intra-Amniotic Infection, PPH: Post-Partum Hemorrhage, HTN: Hypertension, PEC: Preeclampsia SF: Severe Features, PCS: Pelvic Congestion Syndrome, NI: Class I No signs or symptoms, Di/Di: Dichorionic/Diamniotic, D&C: Dilation & Curettage, GDM: Gestational Diabetes Mellitus, PROM: Premature Rupture of Membranes, PTL: Pret-term Labor, PAPP-A: Pregnancy-associated Plasma Protein A, UCTD: Undifferentiated Connective Tissue Disorder, PIH: Pregnancy-Induced/Gestational Hypertension, HCV: Hepatitis C Virus+, ITP: Immune Thrombocytic Purpura

|  | Mother Clinical Presentation |  |  |  |  | Fetal Outcome |  |  | Pregnancy Pathologies |  |  |  |
| --- | --- | --- | --- | --- | --- | --- | --- | --- | --- | --- | --- | --- |
|  | Case | Mat Age | Gest Age | Mother COVID +/− | Patient History | Birth weight | Apgar 1 min | Apgar 5 min | FVM | MVM | Other pathology | Placenta Viral RNA |
| High Positive | P1 | 35 | 25 | + | Symptomatic COVID-19, DFM, Delivered due to nonreassuring fetal status | 650 | 1 | 8 | - | - | MCI | + |
|  | P2 | 34 | 30 | + | Symptomatic COVID-19, DFM, IUFD | 1389 | 0 | 0 | - | - | MFI, CHI | + |
| Positive | P3 | 29 | 40 | + | Symptomatic COVID-19 | 3400 | 9 | 9 | + | - | - | + |
|  | P4 | 40 | 36 | + | T2D (poorly controlled), Placenta previa | 2680 | 9 | 9 | - | - | - | + |
|  | P5 | 16 | 32 | + | Symptomatic COVID-19, PTL | 1740 | 9 | 8 | - | + | - | + |
|  | P6 | 26 | 37 | + | Asymptomatic COVID-19, T2D (poorly controlled), DFM, IUFD | 3200 | 0 | 0 | - | - | Villous Dysmaturity | + |
|  | P7 | 31 | 39 | + | Symptomatic COVID-19 | 3140 | 9 | 9 | - | - | Mec, IVT | + |
|  | P8 | 30 | 38 | + | Symptomatic COVID-19 | 3910 | 9 | 9 | + | - | VUE, Mec | + |
|  | P9 | 41 | 39 | + | Asymptomatic COVID-19 | 3770 | 9 | 9 | - | - | VUE | + |
|  | P10 | 28 | 39 | + | Asymptomatic COVID-19 | 3300 | 9 | 9 | - | - | Mec | + |
|  | P11 | 34 | 37 | + | ICP | 2900 | 9 | 9 | - | - | - | + |
|  | P12 | 31 | 40 | + | Asymptomatic COVID-19 | 3340 | 9 | 9 | - | - | Mec | + |
| Borderline (+) | P13 | 30 | 38 | + | Symptomatic COVID-19, Intrapartum chorioamnionitis | 3360 | 9 | 9 | + | - | Mec, Furcate cord | + |
|  | P14 | 26 | 38 | + | Asymptomatic COVID-19 | 3050 | 9 | 9 | + | - | - | + |
|  | P15 | 19 | 38 | + | Symptomatic COVID-19 | 2390 | 9 | 9 | + | + | ACA | + |
|  | P16 | 28 | 40 | + | Asymptomatic COVID-19 | 3820 | 9 | 9 | + | - | Mec | + |
|  | P17 | 28 | 41 | + | Symptomatic COVID-19 | 4020 | 8 | 9 | + | - | VUE, IDA, Mec | + |
|  | P18 | 41 | 40 | + | Symptomatic COVID-19 | 4115 | 9 | 9 | + | - | Mec | + |
|  | P19 | 32 | 38 | + | Asymptomatic COVID-19 | 3160 | 9 | 9 | - | - | Twisted Cord | + |
|  | P20 | 26 | 39 | + | Symptomatic COVID-19 | 3720 | 9 | 9 | + | - | Hofbauer hyperplasia | + |
|  | P21 | 20 | 38 | + | Symptomatic COVID-19 | 3685 | 6 | 9 | + | - | IAI, Mec | + |
|  | P22 | 25 | 39 | + | Asymptomatic COVID-19 | 3000 | 9 | 9 | - | - | Villitis | + |
| Negative Samples | P23 | 40 | 39 | + | Asymptomatic COVID-19 | 3720 | 9 | 9 | + | - | - | - |
|  | P24 | 40 | 37 | + | Symptomatic COVID-19, PEC | 2060 | 8 | 9 | - | + | Mec | - |
|  | P25 | 38 | 39 | + | Symptomatic COVID-19, ITP, Protein S deficiency, Gestational HTN | 3890 | 9 | 9 | + | - | Funisitis | - |
|  | P26 | 26 | 40 | + | Chronic HTN, Symptomatic COVID-19 | 3799 | 9 | 9 | + | - | - | - |
|  | P27 | 37 | 39 | + | Asymptomatic COVID-19, Autoimmune gastritis | 2415 | 9 | 9 | - | + | - | - |
|  | P28 | 40 | 33 | + | Symptomatic COVID-19, IUGR, PEC, Delivered for nonreassuring fetal status | 1690 | 8 | 8 | - | + | Mec | - |
|  | P29 | 33 | 35 | + | Symptomatic COVID-19, Dichorionic twins, PEC | 2280 (A)<br>2810 (B) | 8 (A)<br>8 (B) | 9 (A)<br>9 (B) | + | + | VUE | - |
|  | P30 | 23 | 39 | + | Asymptomatic COVID-19 | 3580 | 8 | 9 | - | - | Villitis | - |
|  | P31 | 25 | 38 | + | Asymptomatic COVID-19 | 3920 | 9 | 9 | - | - | Mec | - |
|  | P32 | 34 | 39 | + | Symptomatic COVID-19 | 3360 | 9 | 9 | - | - | Mec | - |
|  | P33 | 40 | 37 | + | Asymptomatic COVID-19 | 3400 | 8 | 9 | - | - | - | - |
|  | P34 | 37 | 41 | + | Symptomatic COVID-19 | 3900 | 9 | 9 | - | - | IAI, Mec | - |
|  | P35 | 39 | 37 | + | Asymptomatic COVID-19 | 2650 | 9 | 9 | - | - | Villous dysmaturity | - |
|  | P36 | 34 | 39 | + | Asymptomatic COVID-19 | 2510 | 9 | 9 | - | + | - | - |
|  | P37 | 33 | 23 | + | COVID-19 remote from delivery, fetal anencephaly, intrapartum fetal demise | 370 | 0 | 0 | - | - | - | - |
|  | P38 | 30 | 39 | + | Asymptomatic COVID-19 | 3910 | 9 | 9 | + | - | Villitis | - |
|  | P39 | 31 | 40 | + | Symptomatic COVID-19 remote from delivery | 3200 | 8 | 9 | - | - | - | - |
|  | P40 | 30 | 39 | + | Symptomatic COVID-19 remote from delivery | 3650 | 9 | 9 | + | - | Chorionic cysts, IAI, Mec | - |
|  | P41 | 27 | 41 | + | Symptomatic COVID-19, Low PAPP-A, Gestational HTN | 3630 | 8 | 9 | - | - | Focal chorangiosis | - |
|  | P42 | 23 | 37 | + | Symptomatic COVID-19 remote from delivery | 2510 | 9 | 9 | - | - | IVT | - |
|  | P43 | 31 | 36 | + | ICP, Asymptomatic COVID-19 | 3290 | 9 | 9 | - | - | IVT, Mec | - |
|  | P44 | 29 | 37 | + | Symptomatic COVID-19, low PAPP-A | 2350 | 8 | 9 | - | + | - | - |
|  | P45 | 29 | 36 | + | Genetic carrier for hearing loss | 2510 | 9 | 9 | - | - | - | - |
|  | P46 | 40 | 38 | + | Symptomatic COVID-19 remote from delivery | 2820 | 9 | 9 | - | + | - | - |
|  | P47 | 32 | 40 | + | Symptomatic COVID-19 remote from delivery | 3360 | 9 | 9 | - | + | VUE | - |
|  | P48 | 51 | 37 | + | Symptomatic COVID-19 remote from delivery, T2D | 3080 | 9 | 10 | - | - | VUE | - |
|  | P49 | 41 | 38 | + | Symptomatic COVID-19 remote from delivery, Asthma | 2990 | 8 | 9 | + | + | - | - |
|  | P50 | 38 | 39 | + | Symptomatic COVID-19 remote from delivery | 3010 | 9 | 9 | - | - | VUE, IVT | - |
|  | P51 | 38 | 39 | + | Symptomatic COVID-19 remote from delivery | 3840 | 9 | 9 | - | - | - | - |
|  | P52 | 33 | 38 | + | Symptomatic COVID-19, Long QT syndrome, remote from delivery | 3005 | 9 | 9 | - | - | - | - |
|  | P53 | 38 | 36 | + | Symptomatic COVID-19 remote from delivery, Dichorionic twins, PTL | 2680 (A)<br>2740 (B) | 9 (A)<br>9 (B) | 9 (A)<br>9 (B) | - | - | - | - |
|  | P54 | 35 | 39 | + | Symptomatic COVID-19 remote from delivery | 2870 | 9 | 9 | + | + | Villitis | - |
| Negative Controls | C1 | 32 | 40 | - | Subglottic stenosis | NA | 9 | 9 | - | - | Mec | - |
|  | C2 | 29 | 39 | - | Low PAPP-A, UCTD, celiac disease | 3470 | 9 | 9 | - | - | Mec | - |
|  | C3 | 39 | 34 | - | PPROM | 2320 | 9 | 9 | - | - | IVT | - |
|  | C4 | 31 | 40 | - | COVID-19 in first trimester | 3392 | 9 | 9 | - | - | IVT | - |
|  | C5 | 36 | 39 | - | Gestational HTN, GDM | 3277 | 9 | 9 | - | - | ACA | - |
| Inflammatory Samples | I1 | 32 | 38 | - | Intrapartum chorioamnionitis | 3145 | 9 | 9 | - | - | ACA, Acute funisitis, Mec | - |
|  | I2 | 28 | 40 | - | No medical history | 3100 | 9 | 9 | - | - | VUE, ACA | - |
|  | I3 | 33 | 38 | - | Opioid use disorder, HCV, Placental abruption | 2664 | 9 | 9 | + | - | VUE, Acute funisitis | - |
|  | I4 | 42 | 38 | - | Gestational HTN, GDM | 2891 | 9 | 9 | - | - | ACA, Acute funisitis | - |
|  | I5 | 34 | 39 | - | ITP | 3447 | 7 | 9 | - | + | ACA, Acute funisitis, Mec | - |

The pregnant women ranged in age from 16 to 51 years, with a majority in their 20’s and 30’s. Two pregnancies resulted in intrauterine fetal demise (IUFD) (P2, P6), and one fetus, delivered preterm at 25 weeks of gestation, was admitted to the neonatal intensive care unit (NICU) where the infant has remained for 4 months (P1). All neonates were tested by nasopharyngeal swabs for SARS-CoV-2 at 24 hours, and none were positive. Among the placentas delivered from mothers who tested positive for SARS-CoV-2, 31% (17 cases) presented fetal vascular malperfusion (FVM), 19% (10 cases) displayed maternal vascular malperfusion (MVM), and 7% (4 cases) overlapped for both placental pathologies. None of the healthy control placentas from SARS-CoV-2 negative mothers displayed FVM or MVM (Table 1).

RNA was isolated from all placental samples and subjected qRT-PCR to determine the presence of genomic and replicating SARS-CoV-2 RNA. 22 out of 54 placentas from SARS-CoV-2 positive mothers showed presence of viral RNA (42%), 2 of those were highly positive (4%), 10 positive (19%) and 10 were borderline positive (19%) (Table 1 and Figure 1A). Presence of SARS-CoV-2 in the placenta did not correlate with observed FVM: Of the 22 positive placentas, 10 displayed FVM while 12 were without FVM (Table 1).

**Figure 1.**
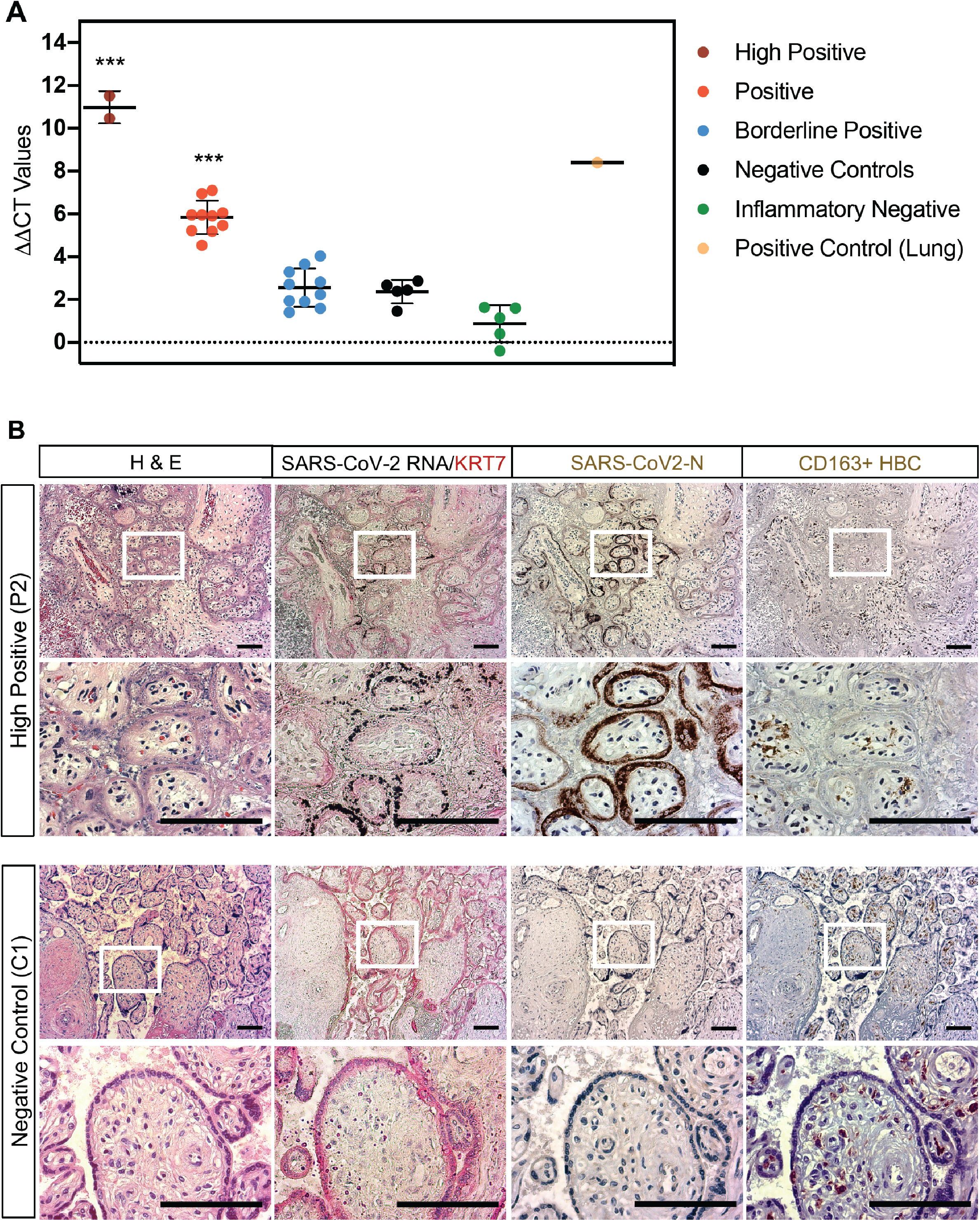
SARS-CoV-2 virus is present in placentas from infected mothers and results in inflammatory responses. (A) Graph showing ΔΔCT values of RNA samples isolated from FFPE patient placenta slides from the different cohorts. A student’s t-test comparing the 3 positive cohorts (High Positive, Positive, Borderline Positive) to the negative control cohort resulted in statistically significant higher viral load in the High Positive and Positive cohorts (*** = p-value < 0.001). (B) Brightfield microscopy images of a representative COVID High positive patient (P3) and a representative negative control patient sample (C1). Slides were stained by H&E, *in situ* PLAYR for SARS-CoV2-RNA counterstained for syncytial trophoblast marker cytokeratin (KRT7, red), and by immunohistochemistry for SARS-CoV2-N protein (brown) as well as for CD163+ Hofbauer cells (HBC). Scale bars = 100μm.

Strikingly, the three pregnancies from SARS-CoV-2 positive mothers that resulted in IUFD or admission of the neonate to the NICU delivered placentas that were highly positive (P1 and P2) or positive (P6) for SARS-CoV-2 (Table 1, grey shaded rows).

### Placental syncytiotrophoblast are the primary target for SARS-CoV-2 infection of pregnant females at term

To determine whether the placenta itself was infected by SARS-CoV-2, qRT-PCR using primers against SARS-CoV-2-N was run on RNA isolated from FFPE patient placenta slides. This provided us with 5 distinct cohorts for this study depicted in Table 1: High Positive (P1, P2; ddCT value > 9), Positive (P3-P12; ddCT > 4.5), and Borderline Positive (P13-P22). We also ran qRT-PCR on RNA from patient placenta samples from SARS-CoV-2 negative mothers (C1-C5) as well as SARS-CoV-2 negative mothers that had unrelated inflammatory pathologies (I1-I5) (Figure 1A and Table 1). To confirm presence of viral RNA, the presence of a distinct amplicon on PCR melt curve and on gels run on RNA samples distinguished between the positive and negative samples from all of the placenta samples obtained from COVID-positive mothers (data not shown).

To identify the cells in the placental chorionic villi that were infected by SARS-CoV-2, adjacent placental sample sections (10 microns apart) from the different cohorts were stained by hematoxylin and eosin (H&E), or for the presence of replicating viral RNA, for the presence of SARS-CoV-2 nucleocapsid protein (SARS-CoV-2-N), and for the presence CD163^+^ Hofbauer cells (HBC) and macrophages using immunohistochemistry. SARS-CoV-2 RNA was detected by *in situ* hybridization in the high positive samples, but not in negative control samples (Figure 1B). Presence of SARS-CoV-2 RNA was restricted to the Keratin-7 (KRT7)-positive syncytial trophoblast layer which anatomically encapsulate the chorionic villi structures (Figure 1B). Similarly, expression of the SARS-CoV-2 N protein was detected in adjacent sections within the same villi. Localization of the N protein was restricted to the syncytiotrophoblast layers in the high positive placentas. Interestingly, the syncytial trophoblast layer of the high positive sample had significantly fewer nuclei and appeared damaged as indicated by the hematoxylin counterstaining. Importantly, at low magnification, massive infiltration of maternal immune cells, including CD163+ macrophages detected in the high positive samples, but not in the controls (Figure 1B and not shown). This result is consistent with the pathology report for the sample P2 of chronic histiocytic intervillositis (CHI) (Table 1). Intravilllous HBC and intervillous maternal macrophages did not show infection with SARS-CoV-2, evidenced by the absence of SARS-CoV-2 RNA and N protein. In summary, these results indicate that syncytiotrophoblast are the primary targets for SARS-CoV-2 infection in the placenta, and that the massive maternal immune cell migration occurred in response to the SARS-CoV-2 infection either in the mother or in the placenta.

### Placental explant and cell cluster cultures are permissive to pseudo-entry virus and infection can be blocked by anti-ACE2 and anti-NRP1 antibodies

To determine the SARS-CoV-2 tropism and infection of term placentas, we used fresh placental isolates from SARS-CoV-2 negative mothers obtained immediately post-delivery. After removal of the fetal chorionic plate and maternal decidua, samples containing terminal, intermediate and stem chorionic villi were used for the preparation of placental villi explant cultures (Figure 2C). In addition, placental cell clusters were prepared by enzymatic digestion of the chorionic villi, followed by filtration that allows passage of small cell clusters. Placental cultures were infected with a dual nanoluciferase/green fluorescent protein (GFP) reporter lentiviral vector pseudotyped with SARS-CoV-2 spike (S) (Tada et al. 2020), and luciferase activity was quantified 72 hpi. Lentiviral reporter viruses pseudotyped with vesicular stomatitis virus G protein (VSV-G) were used as a positive control for infection (Figure 2). A comparison of the infectivity showed that both pseudotyped viruses infected the placental cultures at similar levels. Explant cultures consistently showed an approximately 5-fold lower infection as compared to clusters or single cells, likely due to the reduced surface accessibility. Infectivity was significantly reduced by adding the human immunodeficiency virus (HIV) reverse transcriptase (RT) inhibitor nevirapine (NVP), indicating that the luciferase activity was primarily due to viral entry and not carry-over from residual viral particles in the cultures (Figure 2A). The major SARS-CoV-2 entry factors, ACE2 and TMPRSS2 are expressed in the placenta, albeit their expression is significantly decreased in the third trimester (Pique-Regi et al. 2020; Singh, Bansal, and Feschotte 2020; Ouyang et al. 2021). Furthermore, NRP1 has been identified as a novel host factor for SARS-CoV-2 (Cantuti-Castelvetri et al. 2020; Daly et al. 2020) and is expressed on syncytiotrophoblast (Arad et al. 2017; Baston-Buest et al. 2011). To determine if ACE2 and/or NRP1 facilitate infection in the placenta, we pre-treated placental cell clusters with anti-ACE2 or anti-NRP1 blocking antibody prior to infection. Both antibodies reduced infectivity of SARS-CoV-2 S protein pseudotyped lentivirus in placental cell clusters by about 50%, while anti-ACE2 blocking antibody did not reduce infectivity of VSV-G pseudotyped lentivirus. Pre-treatment with both antibodies did not result in further reduction of infectivity, suggesting the possibility of alternative receptor(s) (Figure 2B).

**Figure 2.**
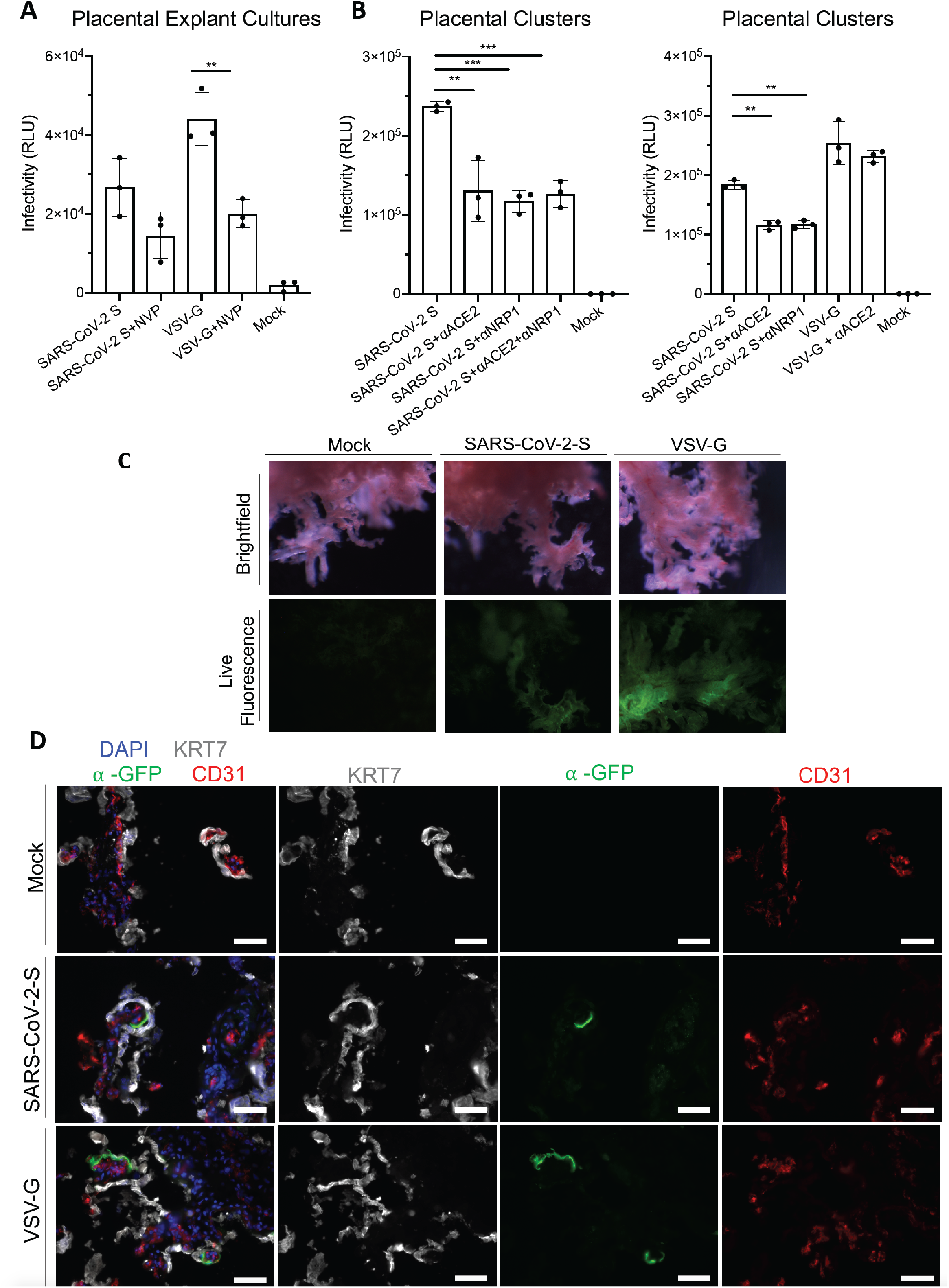
Placental explant and cell clusters can be infected by SARS-CoV-2 S protein pseudotyped lentivirus and infection can be blocked by anti-ACE2 and anti-NRP1 antibodies. (A) Graphs showing relative luminescence units (RLU) from infected explant cultures 72 hpi with the addition of reverse transcriptase inhibitor, Nevirapine (NVP). (B) Graphs showing RLU from infected isolated primary placental clusters 72 hpi with the addition of blocking antibodies against ACE2, NRP1. Statistical analysis was performed using one-way Anova (** = p-value < 0.005, *** = p-value < 0.001). (C) Brightfield and live fluorescence microscopy images of cultured placental explants Mock (left column), or 72hpi with either Lenti-SARS-CoV2-S Pseudovirus (center column) or Lenti-VSV-G (right column). (D) Fluorescence microscopy on mock (top row) and Lenti-SARS-CoV2-S infected (center row) or Lenti-VSV-G infected (bottom row) explant sections stained for the GFP reporter (green) syncytial trophoblast marker, cytokeratin (KRT7, grey), endothelial marker CD31 (red) and DAPI nuclear stain (blue). Scale bars = 500μm.

To determine which cells are targeted by the pseudotyped virus, placental explant cultures were infected for 72 hours with lentivirus pseudotyped by SARS-CoV-2 spike (S) protein, and live GFP could be visualized in the infected explant cultures, with a more robust signal observed in the pseudotyped VSV-G infected cultures (Figure 2C). Explant cultures were then processed for immunofluorescence staining and analyzed for co-localization of the GFP reporter with KRT-7/Cytokeratin (trophoblast marker) and CD31 (endothelial marker). GFP was detected in small patches of syncytiotrophoblast cells located on the outer perimeter of the chorionic villi, but not in endothelial cells (Figure 2D). Similar results were obtained after infection with VSV-G pseudotyped lentivirus, with more robust infection visualized by live fluorescence microscopy on the infected explant cultures, whereas no GFP signal was found in mock-infected explant cultures (Figure 2D).

### Primary placental cell clusters are permissive to SARS-CoV-2

To further determine the intrinsic susceptibility of placental cells to SARS-CoV-2, primary placental cell clusters were infected *ex vivo*. Placentas were isolated from healthy term deliveries as described above, digested into cell clusters of approximately 50-100 cells and plated on Matrigel-coated plates. Cell clusters were infected with live SARS-CoV-2 virus (Isolate USA-WA1/2020, multiplicity of infection, MOI=1). Cells were collected 24 hpi and virus load analyzed by qRT-PCR and immunofluorescence staining.

qRT-PCR analysis using primers targeting subgenomic N transcripts demonstrated robust SARS-CoV-2 viral replication in primary human placental cell clusters at 24 hpi (Figure 3A). To determine which cells of primary placental cell clusters were susceptible to SARS-CoV-2 infection, infected cell clusters were immunostained for SARS-CoV-2 nucleocapsid protein (SARS-N) and cell type specific markers for trophoblast cells (KRT7) and endothelial cells (CD31). Three-dimensional reconstruction of confocal imaging confirmed the presence of SARS-CoV-2-N protein in KRT7+ syncytiotrophoblast (Figure 3B). Co-localization of SARS-CoV-2-N protein was found in multiple clusters of KRT7+ syncytiotrophoblast cells (Figure 3C). In addition to the positive staining for SARS-CoV-2-N in syncytiotrophoblast, there were rare CD31+ endothelial cells that also stained positively for SARS-CoV-2-N protein (Figure 3C).

**Figure 3.**
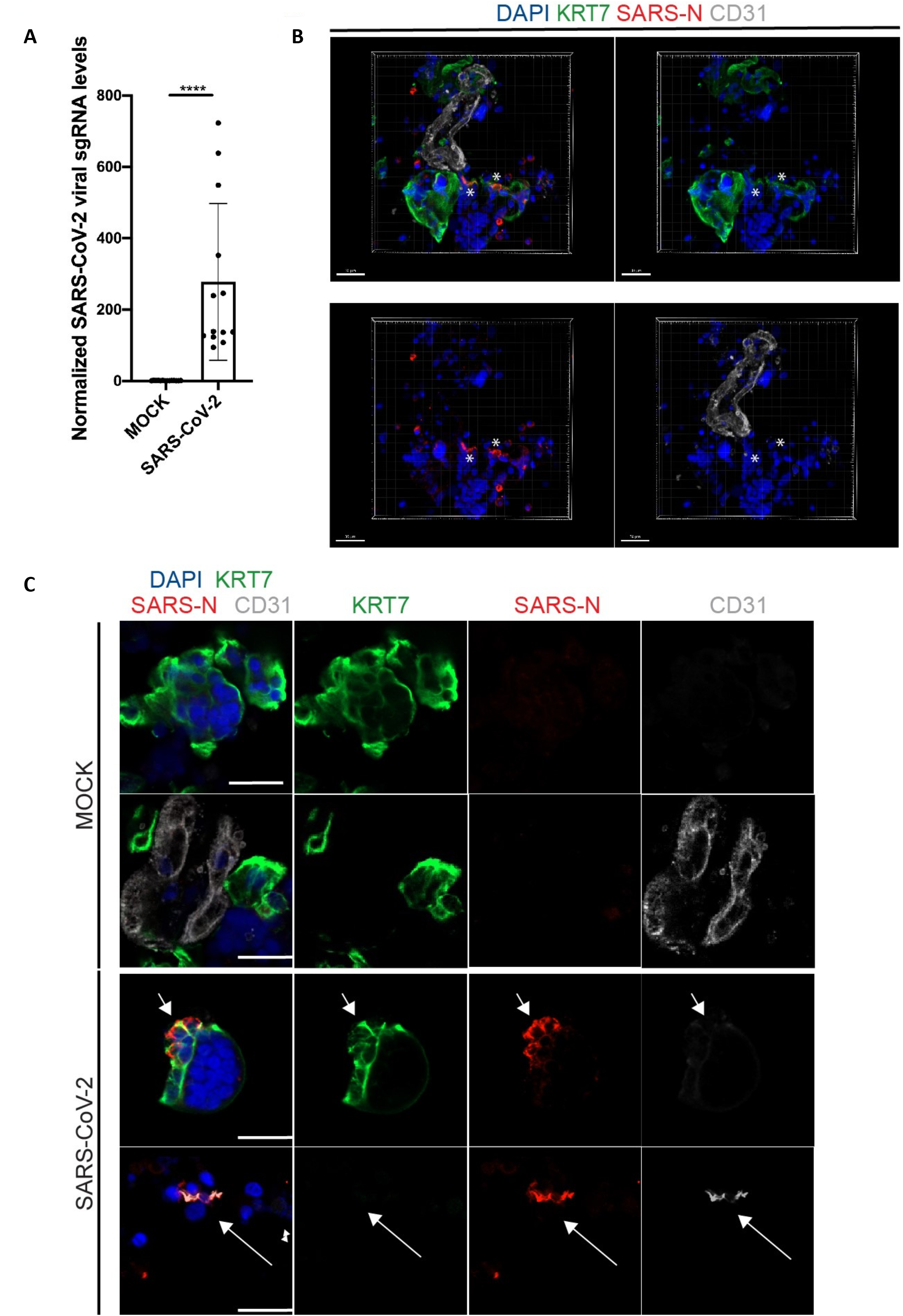
Primary human placenta cells can be infected with SARS-CoV-2 *ex vivo*. (A) qRT-PCR analysis of relative viral N subgenomic RNA expression in primary placental cell clusters infected with SARS-CoV-2 *ex vivo* (MOI=1) for 24 hours and normalized to ACTB levels. (mean+/− SD; n=12 from 4 repeated experiments; ****p<0.0001). (B) Three-dimensional reconstruction of confocal imaging of primary placental cell clusters infected with SARS-CoV-2 *ex vivo* (MOI=1) at 24hpi, stained for trophoblast marker KRT7 (green), SARS-N (red), endothelial marker CD31 (grey), and DAPI (blue). Scale bar = 30μm. (C) Confocal imaging of primary placental cell clusters infected with MOCK (top rows) or SARS-CoV-2 (MOI=1, bottom rows) *ex vivo* at 24hpi, stained for trophoblast marker KRT7 (green), SARS-N (red), endothelial marker CD31 (grey), and DAPI (blue). Arrows indicate presence of SARS-N nucleocapsid protein in trophoblast and endothelial cells. Scale bar = 20μm.

## Discussion

The placenta is a vital organ that provides the gestational interface between mother and fetus. Compromised maternal health and environmental insults, such as viral infections, can result in placental dysfunction and lead to pregnancy complications with increased morbidity and mortality for the mother and fetus (Rossant and Cross 2001; Maltepe, Bakardjiev, and Fisher 2010; John and Hemberger 2012). Impairment of placental function can also developmentally program the fetus for chronic disease later in life, including cardiovascular disease, diabetes and obesity (Burton, Fowden, and Thornburg 2016).

The aim of our study was to investigate the impact of late pregnancy SARS-CoV-2 infection on maternal and fetal health and proper placental function. Within a cohort of 54 placental samples from women who tested positive for SARS-CoV-2 at delivery, 22 were positive for genomic and replicating viral RNA. The observed percentage of positive samples was higher compared to other studies (Hecht et al. 2020b; Facchetti et al. 2020; Debelenko et al. 2021; Lu-Culligan et al. 2021) and likely reflects the fact that New York City was at the epicenter for COVID-16 in March-May of 2020. Furthermore, many of the placental samples were obtained from deliveries where the mothers or infants presented with clinical pathologies which may bias sample collection. Quantification of virus content in the placental samples revealed that the 2 cases with high SARS-CoV-2 presence in the placenta were from pregnancies with adverse fetal outcome, including fetal demise and preterm delivery. In contrast, only 1 out of 10 pregnancies with medium viral content resulted in IUFD, and this may have been triggered by poorly controlled maternal T2D. Of the remaining pregnancies with medium or low virus content in the placenta, all babies tested negative for SARS-CoV-2 and were healthy at discharge. It will be important to follow up on the health of these infants to investigate possible long-term effects of SARS-CoV-2. None of the pregnancies of SARS-CoV-2 negative healthy controls (n=5) or inflammation controls (n=5) resulted in fetal demise.

Upon examining sections from placentas with high virus content, we detected SARS-CoV-2 RNA and protein in a large fraction of syncytiotrophoblast, the single cell layer enveloping the fetal chorionic villi situated at the interphase to maternal blood. No virus was detected in fetal macrophages (Hofbauer cells), other cell types inside the villi, including stromal and endothelial cells, or outside the villi. Several recent reports also provided evidence for SARS-CoV-2 infection restricted to syncytiotrophoblast of placentas from SARS-CoV-2 positive mothers (Alamar et al. 2020; Mulvey et al. 2020; Hecht et al. 2020b; Penfield et al. 2020; Hosier et al. 2020; Vivanti et al. 2020; Taglauer et al. 2020; Facchetti et al. 2020). However, two reports on a preterm placenta and a placenta from a newborn with vertically SARS-CoV-2 noted also presence of SARS-CoV-2 in other cell types, including Hofbauer cells and stromal cells inside the villi, and maternal macrophages and epithelial cells at the maternal-fetal interface (Verma et al., 2021, Fachetti et al, 2020). It is possible that SARS-CoV-2 infection in these cases occurred at earlier gestational stages and allowed for additional viral spread beyond the syncytiotrophoblast layer.

Placentas in our study with high virus presence also displayed massive infiltration of maternal immune cells, including macrophages into the intervillous space. However, we did not detect SARS-CoV-2 in the infiltrating immune cells. Several recent studies reported similar findings; interestingly they were most prominently found in placentas from live-borne and stillborn neonates that had tested positive for SARS-CoV-2. These finding included intervillous infiltration by inflammatory immune cells, chronic histiocytic intravillositis with trophoblast necrosis, and increased fibrin deposition (Debelenko et al. 2021; Facchetti et al. 2020; Garrido-Pontnou et al. 2021; Lu-Culligan et al. 2021; Schwartz et al. 2021; Morotti et al. 2021; Verma et al. 2021). Furthermore, transcriptome data presented in one of these studies showed localized inflammatory responses to systemic SARS-CoV-2 infection in the placenta, even if SARS-CoV-2 virus was not detected (Lu-Culligan et al. 2021).

Considering the low numbers of placental infections by SARS-CoV-2 so far seen clinically, we decided to complement our *in vivo* studies by using *ex vivo* placental explant and cell cluster culture models to study virus entry. We showed infection with SARS-CoV-2 virus or SARS-CoV-2 spike S pseudotyped lentivirus targeted predominantly syncytiotrophoblast and, in rare instances, endothelial cells. Term placentas express very low levels of the SARS-CoV-2 receptor ACE2 and the co-factor TMPRSS2 (Pique-Regi et al. 2020; Singh, Bansal, and Feschotte 2020; Ouyang et al. 2021). Recently, Lu-Culligan et al. (Lu-Culligan et al. 2021) reported on increased levels of placental ACE2 expression in COVID-19 positive mothers; whereas a second publication reported on a decrease of ACE2 expression and dysregulation of the renin-angiotensin system (Verma et al. 2021). We asked here if alternative entry factors might be used by SARS-CoV-2 to infect placental cells. One likely candidate is the transmembrane protein NRP1, which has recently been identified as a host factor that facilitates SARS-CoV-2 cell entry and infectivity (Cantuti-Castelvetri et al. 2020; Daly et al. 2020). NRP1 was originally identified as a co-receptor for Vascular endothelial growth factor (VEGF) on endothelial cells but is expressed also at the maternal-fetal interface in decidual cells and syncytiotrophoblast, and is thought to play important roles during pregnancy and in the immune system (Arad et al. 2017; Baston-Buest et al. 2011). We show that infection by SARS-CoV-2 S pseudotyped lentivirus can be partially inhibited by using blocking anti-ACE2 or anti-NRP1 antibodies, whereas infection by VSV-G pseudotyped lentivirus is not blocked, suggesting that SARS-CoV-2 may use NRP1 as an alternative entry factor, and suggesting the existence of additional entry factors in the placenta.

The present study focused on late cohort infections from mothers who tested positive at the time of delivery. It will be important to study the impact of early cohort infections in mothers who are serologically positive at delivery but negative for viral RNA, as infection during the first and second trimester may affect placental development and morphogenesis and result in different placental pathologies and clinical outcomes for mother and fetus.

## Materials and Methods

### Cell Lines

Vero E6 (African green monkey [Chlorocebus aethiops] kidney) were obtained from ATCC (https://www.atcc.org/). Vero E6 and A549 (adenocarcinomic human alveolar basal epithelial cell line)-ACE2 cells were cultured in Dulbecco’s Modified Eagle Medium (DMEM) supplemented with 10% fetal bovine serum (FBS) and 100 U/mL penicillin and 100 μg/mL streptomycin, and maintained at 37°C with 5% CO_2_.

### SARS-CoV-2 propagation and infection

SARS-CoV-2 isolate USA-WA1/2020 (NR-52281) was provided by the Center for Disease Control and Prevention (CDC) and obtained through BEI Resources, NIAID, NIH. SARS-CoV-2 was propagated in Vero E6 cells in DMEM supplemented with 2% FBS, 4.5 g/L D-glucose, 4 mM L-glutamine, 10 mM Non-essential amino acids, 1 mM sodium pyruvate and 10 mM HEPES using a passage-2 stock of virus. Three days after infection, supernatant containing propagated virus was filtered through an Amicon Ultra 15 (100 kDa) centrifugal filter (Millipore Sigma) at ~4000 rpm for 20 minutes. Flow-through was discarded and virus was resuspended in DMEM supplemented as described above. Infectious titers of SARS-CoV-2 were determined by plaque assay in Vero E6 cells in Minimum Essential Media supplemented with 2% FBS, 4 mM L-glutamine, 0.2% BSA, 10 mM HEPES and 0.12% NaHCO_3_ and 0.7% agar. All MOI values were based on titer determined from plaque assays on Vero E6 cells. All work involving live SARS-CoV-2 was performed in the CDC/USDA-approved biosafety level-3 (BSL-3) facility of the Icahn School of Medicine at Mount Sinai in accordance with institutional biosafety requirements.

### Placental Samples

Placental tissues from SARS-CoV-2 positive women and controls were obtained at delivery at Weill Cornell-NY Presbyterian by the Department of Pathology and Laboratory Medicine at Weill Cornell Medicine. All women admitted for delivery were tested by nasal swabs for acute SARS-CoV-2 infection by qRT-PCR, and serologically for previous infection at Weill Cornell Medicine Department of Pathology and Laboratory Medicine. Infants were tested for SARS-CoV-2 at birth and 1 week of age by nasal swabs and RT-PCR. Placental samples were fixed for 48 hours in formalin and then processed and embedded into formalin fixed paraffin embedded (FFPE) blocks by the pathology department. FFPE placenta samples from 5 healthy women who tested negative for SARS-CoV-2 served as controls. An additional 5 FFPE placental samples with inflammation pathologies, obtained from SARS-CoV-2 negative patients, were also included in the study. Unstained sections and H&E sections of the FFPE blocks were performed at the Weill Cornell Clinical & Translational Science Center (CTSC) core facility. Additional H&E staining was performed by the Weill Cornell Histology core facility.

### SARS-CoV-2 Detection in RNA from FFPE Placental Sections by qRT-PCR

Total RNA samples were prepared from FFPE placental tissue sections, followed by DNaseI treatment using manufacturer’s instructions (Qiagen RNeasy FFPE kit Cat#73604). To quantify viral replication, as measured by the expression of nucleocapsid sub genomic viral RNA along with the housekeeping gene GAPDH, two-step RT-qPCR was performed using LunaScript^®^ RT SuperMix Kit (E3010L) for cDNA synthesis and Luna^®^ Universal qPCR Master Mix (NEB #M3003) for RT-qPCR. Quantitative real-time PCR reactions were performed on CFX384 Touch Real-Time PCR Detection System (BioRad). The sequences of primers/probes are provided below.

SARS-CoV-2-N

Forward 5’ CTCTTGTAGATCTGTTCTCTAAACGAAC 3’

Reverse 5’ GGTCCACCAAACGTAATGCG 3’

GAPDH

Forward 5’ CATCACCATCTTCCAGGAGCGAGAT 3’

Reverse 5’ GAGGCATTGCTGATGATCTTGAGGC 3’

qRT-PCR graphs were generated using GraphPad Prism software.

### RNA *In Situ* Hybridization to Detect SARS-CoV2 RNA on FFPE Placental Sections

#### Probe design

Probes were designed with a 20-25 nucleotides homology to SARS-CoV-2 genomic RNA and were assessed by NCBI BLAST to exclude off target binding to other cellular transcripts. IDT OligoAnalyzer (Integrated DNA Technologies) was used to identify probe pairs with similar thermodynamic properties, melting temperature 45-60°C, GC content of 40-55%, and low self-complementary. The 3’ end of each one of the probes used for proximity ligation signal amplification is designed with a partially complementary sequence to the 61bp long backbone and partially to the 21bp insert as described previously (Yang et al. 2020).

#### Tissue viral RNA staining pretreatment

Sections of FFPE placental samples were deparaffinized using 100% xylenes, 5 min at room temperature, repeated twice. Slides were rinsed in 100% ethanol, 1 min at room temperature, twice and air dried. Endogenous peroxidase activity was quenched by treating the samples with 0.3% hydrogen peroxide, 10 min at room temperature followed by washing with DEPC treated water. Samples were incubated 15 min at 95-100 °C in antigen retrieval solution (ACDBio, Newark, CA, USA) rinsed in DEPC treated water and dehydrated in 100% ethanol, 3 min at room temperature and air dried. Tissue sections were permeabilized 30 min at 40°C using RNAscope protease plus solution (ACDBio, Newark, CA, USA) and rinsed in DEPC treated water.

#### SARS-CoV-2 RNA detection by probes proximity ligation

Hybridization was performed overnight at 40°C in a buffer based on DEPC-treated water containing 2× SSC, 20% formamide (Thermo Fischer Scientific, Waltham, MA, USA), 2.5 % (vol/vol) polyvinylsulfonic acid, 20 mM ribonucleoside vanadyl complex (New England Biolabs, Ipswich, MA, USA), 40 U/ml RNasin (Promega, Madison, WI, USA), 0.1% (vol/vol) Tween 20 (Sigma Aldrich), 100 μg/ml salmon sperm DNA (Thermo Fisher Scientific, Waltham, MA, USA), 100 μg/ml yeast RNA (Thermo Fisher Scientific, Waltham, MA, USA). DNA probes dissolved in DEPC-treated water were added at a final concentration of 100nM (Integrated DNA Technologies, Coralville, IA, ISA). Samples were washed briefly and incubated in 2× SSC, 20% formamide, 40 U/ml RNasin at 40 °C and then washed four times (5 min each) in PBS, 0.1% (vol/vol) Tween 20, and 4 U/ml RNasin (Promega, Madison, WI, USA). Slides were then incubated with 100 nM insert/backbone oligonucleotides in PBS, 1× SSC, 0.1% (vol/vol) Tween 20, 100 μg/ml salmon sperm DNA (Thermo Fisher Scientific, Waltham, MA, USA), 100 μg/ml yeast RNA (Thermo Fisher Scientific, Waltham, MA, USA), 40 U/ml RNasin at 37 °C. After four washes, tissues were incubated at 37°C with 0.1 U/μl T4 DNA ligase (New England Biolabs, Ipswich, MA, USA) in 50mM Tris-HCl, 10mM MgCl_2_, 1mM ATP, 1mM DTT, 250μg/ml BSA, 0.05% Tween 20, 40 U/ml RNasin, followed by incubation with 0.1 U/μl phi29 DNA polymerase in 50 mM Tris–HCl, 10 mM MgCl_2_, 10 mM (NH_4_)_2_SO_4_, 250μM dNTPs, 1mM DTT, 0.05% Tween 20, 40 U/ml RNasin pH 7.5 at 30 °C. Slides were washed and endogenous biotin was blocked using Avidin/Biotin blocking kit (Vector laboratories, Burlingame, CA, USA) according to the manufacture instructions. Rolling cycle amplicons were identified using a biotin labeled DNA probe at a concentration of 5 nM at 37 °C in PBS, 1× SSC, 0.1% Tween 20, 100 μg/ml salmon sperm DNA, 100 μg/ml yeast RNA, 1 hr. After washing, samples were incubated with 1:100 diluted streptavidin-HRP (Thermo Fisher Scientific, Waltham, MA, USA) in PBS, 60 min at room temperature followed by washing. Labeling was accomplished using EnzMet kit (Nanoprobes, Yaphank, NY, USA) according to manufacture instructions. Slides were further labeled with rabbit anti-cytokeratin 1:250 (Dako Z0622), overnight 4°C. After washing, samples were incubated with 1:1000 with anti-rabbit alkaline phosphatase antibody (1:1000, Jackson immunoresearch, Baltimore, PA, USA) and stained using Fast Red substrate kit according the manufacture instructions (Abcam, Cambridge, MA, USA). Hematoxylin was used for counterstaining (Vector laboratories, Burlingame, CA, USA), and samples were mounted in Permount (Fischer Scientific, Waltham, MA, USA).

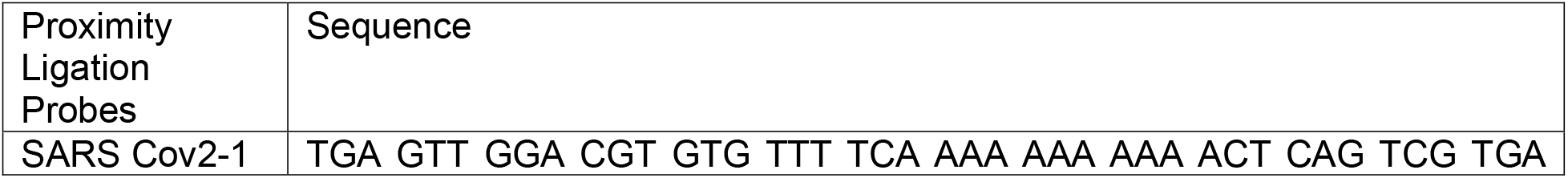
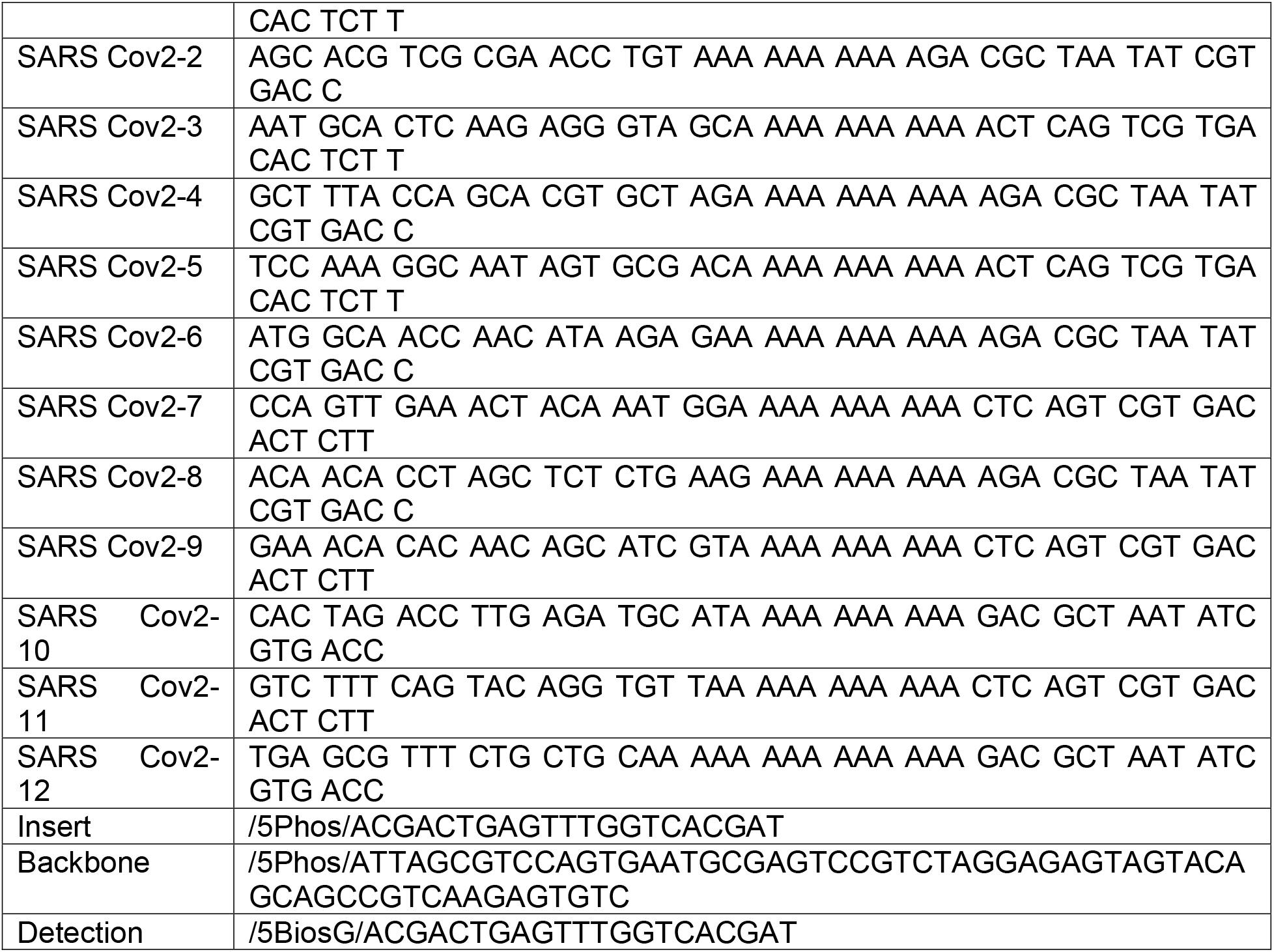

### Placental Explants and Cluster Cultures

Fresh de-identified placentas from SARS-CoV-2-negative mothers were collected within 30 min to 2 hours post-delivery from Labor & Delivery at WCM/NYP. Collection of placentas was performed under an approved IRB exempt protocol (#20-07022453, Weill Cornell Medicine.) Tissue samples were dissected by removing the fetal chorionic plate and any remaining maternal decidual tissue. Primary explant cultures (1cm x 1cm x 2cm) containing terminal, intermediate and stem chorionic villi were further dissected, washed in ice cold 1X PBS to remove maternal blood, and plated into 48-well plastic dishes in DMEM/F12 medium supplemented with 10% FBS and 100 U/mL penicillin, 100 μg/mL streptomycin and 0.25 μg/mL amphotericin B, as previously described (Massimiani et al. 2019).

In addition, placental cell clusters were prepared from fresh chorionic villi tissue samples by mincing with scissors and 10 blade scalpels. The minced tissue was digested using 0.2 mg/mL collagenase/ 0.8U/mL dispase (Roche) and recombinant DNAse I (Sigma) in MACS buffer (PBS/2mM EDTA/ 0.5% bovine serum albumin (BSA)) at 42°C with agitation by pipetting with a 5 ml stripette. The digested tissue was filtered through 100 μ filters (Corning 352360), and red blood cells (RBC) were removed using RBC Lysis Buffer (Biolegend 420301). Clusters were washed once in MACS buffer, examined for viability with Trypan Blue (GIBCO) and plated onto Matrigel-coated 96-well dishes and μ-slide 8-well chamber slides (ibidi GmbH, Germany) at confluent density in DMEM/F12 supplemented with 10% FBS and penicillin/streptomycin/ fungizone, and were incubated at 37°C with 5% CO_2_ for 24 hours prior to infection with pseudovirus to allow for attachment,

For infection with SARS-CoV-2, Sections of fresh chorionic villi (2g) were minced with sterile scalpels, digested in Accutase (Innovative Cell Technologies) for 7 min or isolated using a human umbilical cord dissociation kit (Millitenyi Biotec 130-105-737), and filtrated through a 100 μm cell strainer (Falcon) to obtain cell clusters of ~50-100 cells. Red blood cells were lysed using RBC Lysis Buffer (Biolegend), washed with PBS-0.5% BSA, and resuspended in medium (DMEM-10%FBS-1% Pen-Strep-Glutamax). Cell viability was determined with Trypan blue (Gibco). Cell clusters were plated on Matrigel (Corning, hESC-qualified)-coated plates at 4×10^5/well in 24-well plates or 3×10^4/well in glass-like polymer bottom 96-well plates (CellVis).

### Infection of Ex Vivo Placental Cultures

#### Infection of Explants and Placental clusters with Pseudovirus

Lentiviruses encoding dual Nanoluciferase/GFP reporter lentivirus and pseudotyped by SARS-CoV-2 spike (S) protein (D614G) or VSV-G were prepared as previously described (Tada et al. 2020). The viruses were concentrated 10-fold by ultracentrifugation and titers were quantified by reverse transcriptase assay. Placental explant cultures and cell clusters were infected with 10 μl SARS-CoV-2 S or VSV-G pseudotyped lentivirus (Tada et al. 2020). To determine whether NRP1 or ACE2 facilitates the infection of placental cells, placental cell clusters were pretreated for 30 min with anti-NRP1 mAb (R&D Systems, AF3870) or anti-ACE2 mAb (Agilent, AG-20A-0032-C50). Infected placental clusters were lysed 72 hours post-infection. Luciferase activity was measured using a Promega Nano-Glo Assay Kit and read on an Envision microplate luminometer (Perkin Elmer).

#### Infection of Placental Clusters with Live SARS-CoV-2

Placental cell clusters were infected with live SARS-CoV-2 (isolate USA-WA1/2020 (NR-52281) at an MOI of 0.1 and 1 or mock-infected at day-1 in culture as recently described (Yang et al. 2020). At the indicated hpi, cells were washed three times with PBS. For RNA analysis cells were lysed in TRIzol (Invitrogen). For immunofluorescence staining cells were fixed in 4% formaldehyde for 60 min at room temperature. All work involving live SARS-CoV-2 was performed in the CDC/USDA-approved BSL-3 facility of the Icahn School of Medicine at Mount Sinai in accordance with institutional biosafety requirements.

#### qRT-PCR for Viral Load of SARS-CoV-2 Infected Placental Clusters

Total RNA was extracted using Trizol (Thermo Fisher Scientific, Waltham, MA, USA) followed by ezDNAse treatment (Thermo Fisher Scientific, Waltham, MA, USA) per manufacturer’s instructions. To quantify viral replication, measured by the accumulation of subgenomic N transcripts, one-step quantitative real-time PCR was performed using the SuperScript III Platinum SYBR Green One-Step qRT–PCR Kit (Invitrogen) with primers specific for TRS (listed above) and beta-actin (ACTB) as an internal reference (listed below), as previously described (Yang et al. 2020). Reactions were performed on a QuantStudio 6 Flex Real Time PCR Instrument (Applied Biosystems). The delta-delta-cycle threshold (ΔΔCT) was determined relative to ACTB and mock-infected samples. Graphs were generated using GraphPad Prism software.

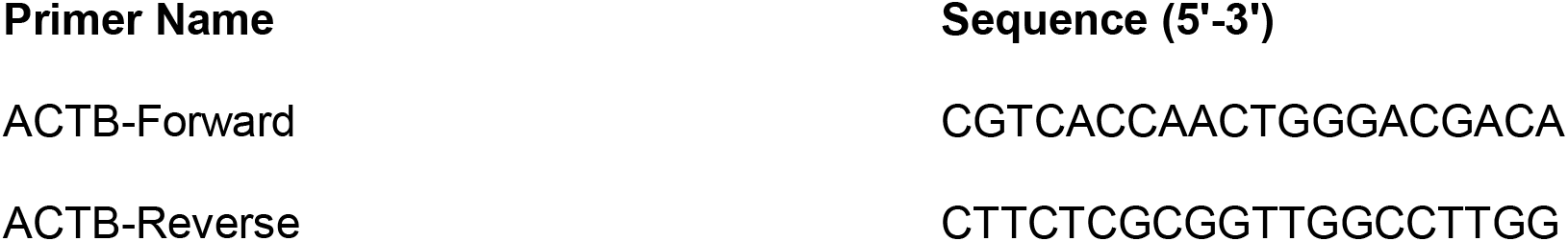

### Immunostaining of FFPE Placental Sections and Infected Placental Cell Clusters

#### IHC for SARS-CoV2 and Hofbauer Cells on FFPE Slides

Immunohistochemistry (IHC) was performed on FFPE slides using ImmPRESS Reagent kit (Vector laboratories, Burlingame, CA, USA). FFPE slides were dewaxed in a hybrid oven for 45 minutes at 55°C and then rehydrated using xylenes followed by a standard ethanol gradient. Antigen retrieval was performed using sodium citrate buffer, pH 6.1 in a steamer for 35 minutes. Slides were blocked using 2.5% horse serum (Vector laboratories) for 1 hour at room temperature and then incubated overnight at 4°C in a humid chamber with primary antibodies (SARS-CoV-2-N, GeneTex GTX635679, at 1:100; CD163,Novus Biologicals NBP2-48846, 1:250) diluted in 1% BSA/0.1% Triton-X PBS (PBST). Slides were treated with 3% hydrogen peroxide at room temperature (Sigma H1009), washed 3 times with 0.1% PBST and then incubated for 1 hour at room temperature with ImmPRESS anti-rabbit peroxidase conjugated antibody (Vector Laboratories, Burlingame, CA, USA). Slides were again washed 3 times with 0.1% PBST with final wash in PBS prior to developing using freshly prepared DAB substrate (Vector Labs SK-4100). Slides were rinsed with water and counterstained with Hematoxylin (RICCA Chemical Company, Arlington, TX, USA) to mark the nuclei. Stained slides were dehydrated using an increasing ethanol gradient, treated with xylenes, and then mounted with Permount solution (Thermo Fisher Scientific, Waltham, MA, USA). Brightfield images were acquired using a Zeiss microscope (Carl Zeiss, Germany).

#### IF Staining for Pseudovirus Infected Placental Explants/Clusters

For immunofluorescence (IF) analysis, SARS-CoV-2 GFP-pseudotyped virus infected explant cultures were drop-fixed overnight in 4% paraformaldehyde (PFA) in PBS containing Ca2^+^/Mg^2+^ at 4°C on a rocker 72 hours post-infection. The fixed explants were then dehydrated with 30% sucrose in PBS overnight at 4°C on a rocker. Explants were embedded in optimal cutting temperature compound (OCT) on dry ice. Frozen blocks were sectioned on a cryomicrotome at 10 micron thickness. Explant culture sections were blocked for 1 hour in 10% donkey serum (Jackson ImmunoResearch labs, Westgrove, PA) in 0.1% PBST. Primary antibodies (rabbit anti-cytokeratin 1:1000 (Dako Z0622), sheep anti-human CD31 1:500 (BD AF806) and chicken anti-GFP 1:1000 (Abcam ab13970)) were diluted in 10% donkey serum-0.1% PBST and incubated overnight at 4°C followed by incubation with secondary antibodies (AlexaFlour647-donkey anti-rabbit, AlexaFlour594-donkey anti-sheep, and AlexaFlour488-donkey anti-chicken, Jackson ImmunoResearch labs, Westgrove, PA)). The clusters were then stained using 4’,6-diamidino-2-phenylindole (DAPI). Slides were mounted with coverslips using ProLong Gold Antifade Mountant with DAPI (Thermo Fisher Scientific, Waltham, MA, USA). Fluorescence microscopy was performed using a Zeiss fluorescent microscope and image analysis was done using ImageJ software.

#### Immunofluorescence Staining for SARS-CoV-2 of Infected Placental Clusters

PFA-fixed cells were blocked in 5% normal donkey serum in PBS-0.05% Triton X-0.01% Saponin (PBS-TSP). Primary antibodies (SARS-CoV2-N, Genetex GTX635679, 1:200; KRT7, Agilent Dako M701829-2, 1:400; PECAM1, R&D AF806, 1:1000) were incubated overnight at 4degC in block, followed by incubation in secondary antibodies (AlexaFluor488-donkey-anti-mouse, AlexaFluor594-donkey-anti-rabbit, AlexaFluor647-donkey-anti-sheep, ThermoFisher, 1:500) in PBS-TSP, and counterstaining with DAPI (Thermo Fisher Scientific, Waltham, MA, USA). Images were acquired using a Zeiss LSM 800 Confocal microscope and processed using Imaris software (Bitplane).

## Acknowledgements

We thank the patients, their families, and healthcare workers fighting the COVID-19 pandemic. This work was supported by a Weill Cornell Medicine COVID-19 Research Grant (H.S., R.E.S., R.N.B. Baergen), the NCI (R01CA234614) and NIAID (2R01AI107301) and NIDDK (R01DK121072) to Department of Medicine, Weill Cornell Medicine (R.E.S.), NIDDK (R01DK119667, R01DK119667-02S1) to S.C. R.E.S. and S.C. are supported as an Irma Hirschl Trust Research Award Scholar. LBA was supported in part by NYSTEM Training grant. L.A.L. is supported by an F32 post-doctoral fellowship from the National Institute of Health (1F32HD096810-01A1) and Weill Cornell Medicine Research Assistance for Primary Parents Award. N.R.L was supported by grants from the NIH (DA046100, AI122390 and AI120898). T.T. was supported by the Vilcek/Goldfarb Fellowship Endowment Fund. We would also like to acknowledge Michael D. Glendenning for his technical assistance with the IHC and the WCM Histology Core.

